# SpatialDWLS: accurate deconvolution of spatial transcriptomic data

**DOI:** 10.1101/2021.02.02.429429

**Authors:** Rui Dong, Guo-Cheng Yuan

## Abstract

Recent development of spatial transcriptomic technologies has made it possible to systematically characterize cellular heterogeneity while preserving spatial information, which greatly enables the investigation of structural organization of a tissue and its impact on modulating cellular behavior. On the other hand, the technology often does not have sufficient resolution to distinguish neighboring cells which may belong to different cell types, therefore it is difficult to identify cell-type distribution directly from the data. To overcome this challenge, we have developed a computational method, called spatialDWLS, to quantitatively estimate the cell-type composition at each spatial location. We benchmarked the performance of spatialDWLS by comparing with a number of existing deconvolution methods using both real and simulated datasets, and we found that spatialDWLS outperformed the other methods in terms of accuracy and speed. By applying spatialDWLS to analyze a human developmental heart dataset, we observed striking spatial-temporal changes of cell-type composition which becomes increasing spatially coherent during development. As such, spatialDWLS provides a valuable computational tool for faithfully extracting biological information from spatial transcriptomic data.

## Main

Rapid development in spatial transcriptomics has enabled systematic characterization of cellular heterogeneity while preserving spatial context^1–6^. Compared to the commonly used singlecell RNA-seq technology, the main advantage of spatial transcriptomic technologies is that they can be used to profile gene expression in a small number of or even single cells while preserving spatial information. This is crucial for mapping the structural organization of tissues and facilitates mechanistic studies of cell-environment interactions. On the other hand, identifying the spatial distributions of various cell types can be challenging, since many existing methods do not have single-cell resolution, such as Spatial Transcriptomics^4^, 10X Genomics Visium, Slide-seq^2^, Dbit-seq^6^, and Nanostring GeoMx. This is an important barrier for data analysis and interpretation which limits the utility of these technologies. Therefore, it is desirable to develop computational methods to infer the composition of cell types at each location, a task that is often referred to as cell-type deconvolution.

A number of methods have been developed for deconvolve bulk RNAseq data^7–13^. In principle, these methods can be directly applied to spatial expression analysis as well, treating the data from each location as a bulk sample. However, there are two main limitations for this approach. First, the number of cells within each spot is typically small. For example, each spot in the 10X Genomics Visium platform has the diameter of 55 μm, corresponding to a spatial resolution of 5-10 cells. The application of a bulk RNAseq deconvolution method to such a small sample size would result in noise from unrelated cell types. Second, as spatial expression datasets usually contain thousands of spots, it would be time and memory consuming if deconvolution methods designed for bulk RNA-seq were applied on spatial expression datasets.

Recently, several methods have been developed specifically for spatial transcriptomics data deconvolution^14–16^. Here, we introduce a novel method spatialDWLS for this task and benchmark with existing methods. In a nutshell, spatialDWLS contains two steps (**Fig. 1a**). First, it identifies cell types that likely to be present at each location by using a recently developed cell-type enrichment analysis method^17^. Second, the cell type composition at each location is inferred by extending the dampened weighted least squares (DWLS) method^9^, which was originally developed for deconvolving bulk RNAseq data. The details are described in the Methods section.

**Figure 1.**
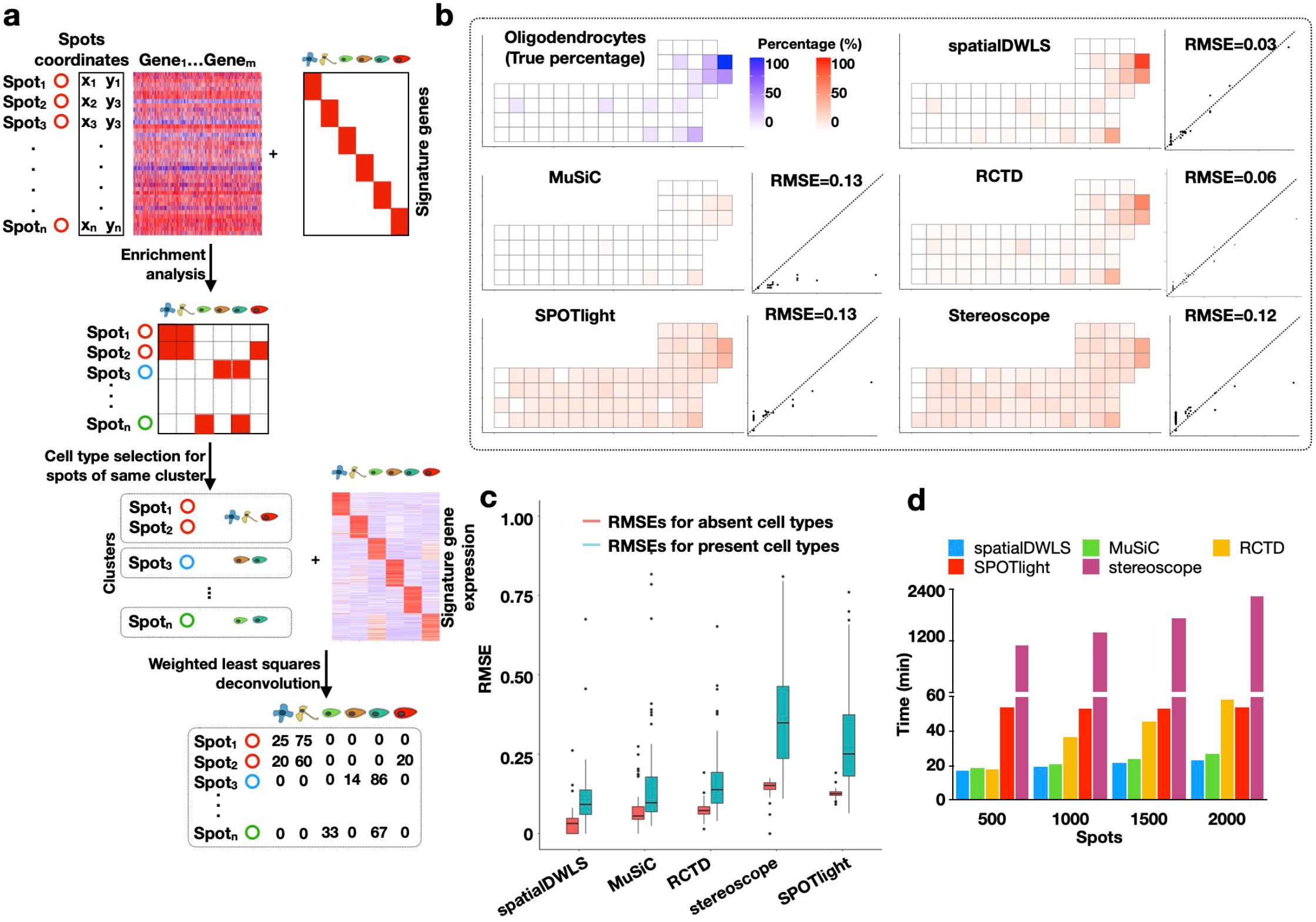
An overview of the spatialDWLS method. **a**. A schematic representation of the spatialDWLS workflow. The input contains a spatial transcriptomic dataset (gene expression matrix and cell location coordinates) and a set of known cell-type specific gene signatures. For each spot, the cell types that are likely to be present are identified by using cell-type enrichment analysis. Then, a modified DWLS method is applied to infer cell type position at each spot. **b.** Comparison of the accuracy of different deconvolution methods. Single-cell resolution seqFISH+ data are coarse-grain averaged to generate lower-resolution spatial transcriptomic data. The true frequency of a cell-type (indicated as blue squares in the top left panel) at each spot is compared with the inferred frequency (indicated as red squares in the five other panels) by using different methods. The relationship is also represented as a scatter plot, with x-axis representing the true frequency and the y-axis representing the inferred frequency. The overall performance is quantified as the root mean square error (RMSE). The oligodendrocyte cell-type is used here as a representative example. **c.** The overall RMSE error is further decomposed into two components, corresponding to regions where the cell type is absent (red) and present (green), respectively. **d.** Comparison of the computing speed of different methods. Running times for analyzing a mouse brain Visium dataset are shown.

To evaluate the performance of spatialDWLS, we created a simulated spatial transcriptomic dataset based on coarse-graining average of single-cell resolution data. Specifically, we analyzed a public seqFISH+ dataset ^1^, which contains the expression profile of 10,000 gene in 523 cells from the mouse somatosensory cortex at the single-cell resolution. To mimic the outcome of a lower-resolution profiling strategy, we divided each field of view (FOV) into squared spots of ~51.5μm on each side, and aggregated the transcript counts that fall into each spot. On average, about seven cells are included in each spot. The resulting dataset has a total number of 71 spots, each covering an average of 7.3 cells. The original dataset serves as the ground-truth for benchmarking.

To apply spatialDWLS, we obtained cell-type specific gene signatures from a publicly available single-cell RNAseq dataset^18^. In total, this dataset contains 1,691 cells and 6 major cell types are identified. Based on the single-cell RNAseq derived cell-type gene signatures, we applied spatialDWLS to deconvolve the above simulated dataset. The cell-type percentage at each location varies from 5.9% to 100%.

To evaluate the performance of our spatialDWLS method, we compared the predicted and true cell type proportion and found good agreement overall (**Fig. 1b-d** and **S1a-b**). For example, the Root Mean Square Error (RMSE) associated with oligodendrocytes is only 0.03 with the predicted values approximately center around ground-truth (**Fig. 1b**). In order to separately evaluate the sensitivity and specificity, we divided the simulated spots into subsets where the cell type is present or absent and evaluated the RMSE errors for each subset. The fact that both errors have small magnitude indicate spatialDWLS has both high degrees of sensitivity and specificity (**Fig. 1c**). As a benchmark, we applied four published deconvolution methods, including MuSiC ^8^, RCTD^14^, SPOTlight^15^ and stereoscope^16^ to analyze the same dataset. All the other methods led to higher error (**Fig. 1b, S1a, b**), although the differences with MuSiC and RCTD appear modest.

Next, we applied spatialDWLS to analyze a 10X Genomics Visium dataset mapping the spatial transcriptomic profile in mouse brain. This dataset contains 2,698 spatially barcoded circular spots each 55μm in diameter. To comprehensively deconvolve cell type composition, we used the mouse nervous system atlas scRNA-seq data as a reference^19^, which contains gene expression signature of 21 major cell types. While it is impossible to quantify the prediction accuracy because the ground-truth is unknown, the resulting spatial distributions are highly consistent with the mouse Allen Brain Atlas (**Fig. S2a, b**). For example, the peptidergic cells were correctly mapped to the hypothalamus region; the granule neurons were correctly mapped to the dentate gyrus region, and the medium spiny neurons were correctly mapped to the basal ganglia (**Fig. S2a, b**).

The spatialDWLS analysis took 23 minutes CPU time on a small computer cluster (Intel Xeon CPU E5-2650 32 processors 2.00GHz and 380Gb memory). To compare the computational efficiency of different methods, we applied each other method to analyze the same dataset using the same computer. Furthermore, to assess scalability we subsampled the mouse brain dataset varying from 500 to 2,000 spots and examined the relationship between CPU time and sample size. We found that spatialDWLS and MuSiC were more computationally efficient, each taking about 23 mins CPU time to analyze the 2000-spot dataset. In comparison, both RCTD and SPOTlight were about 2 times slower for the larger sample size, whereas stereoscope was 10 times slower. Taken together, these analyses suggest spatialDWLS is more accurate and computationally efficient than these other methods.

During embryonic development, the spatial-temporal distribution of cell types changes dramatically. Therefore, it is of interest to test whether spatialDWLS could aid the discovery of such dynamic changes. Recently, Asp and colleagues studied the development of human heart in early embryos (4.5–5, 6.5, and 9 post-conception weeks) by using the Spatial Transcriptomics (ST) technology^20^ (**Fig. 2a**). Since the data does not have single-cell resolution, they were not able to identify cell-type distribution directly from the ST data. In order to apply spatialDWLS, we utilized the single-cell RNAseq derived gene signatures from this study as reference. All the cell types were mapped to expected locations (**Fig. 2b** and **S3a-c**).

**Figure 2.**
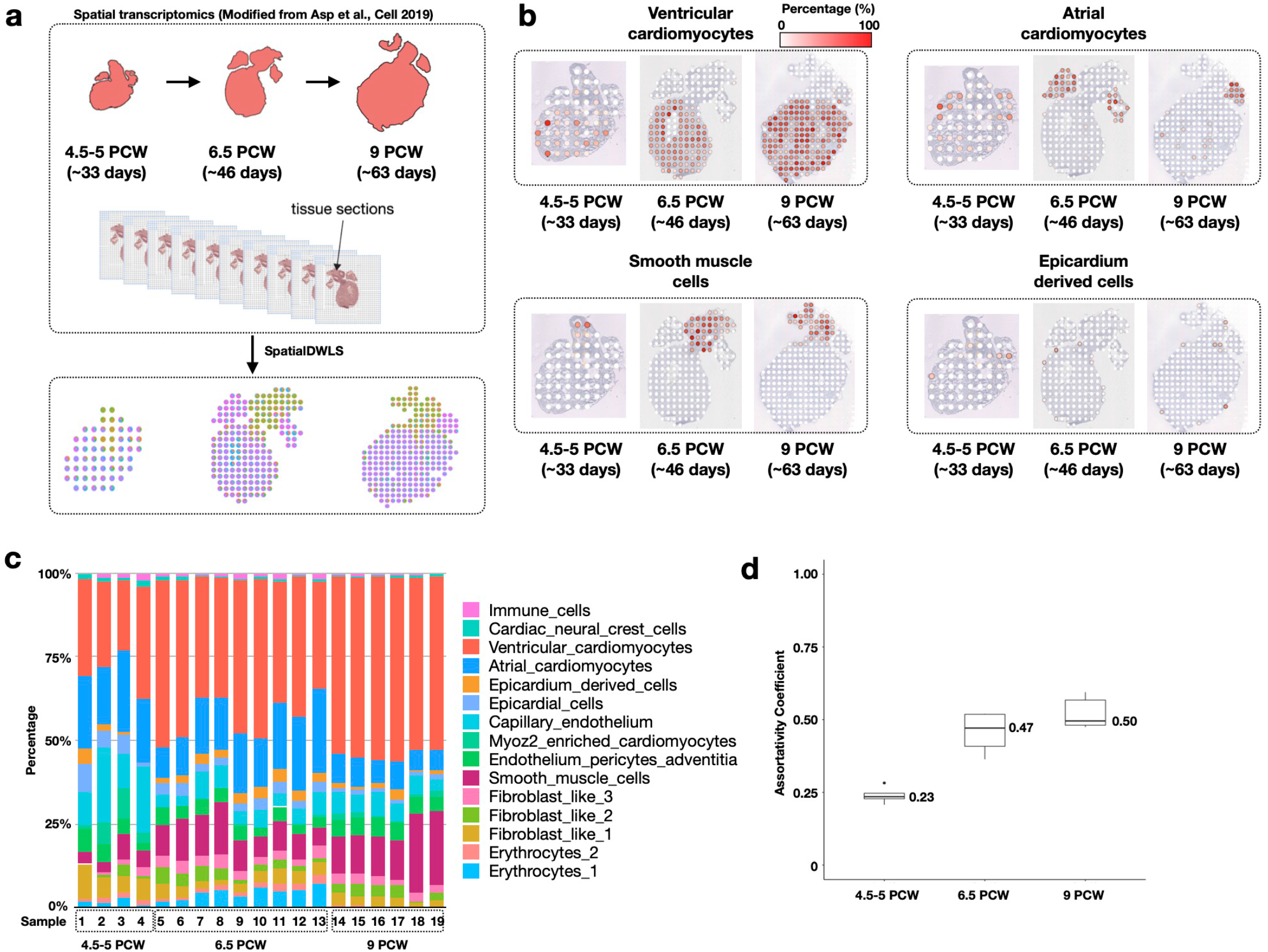
Deconvolution analysis identifies spatial-temporal change of cell-type composition during heart development. **a.** A schematic overview of the analysis. Spatial Transcriptomic data for developing heart were collected at three developmental stages by Asp et al. 2019. In parallel, single-cell RNAseq analysis was carried out to identify cell-type specific gene signatures. The spatialDWLS method was applied to infer the distribution of different cell-types across developmental stages. **b**. The resulting estimates of the spatial distribution of different cell types. One representative sample was selected from each developmental stage. **c**. A summary of the cell-type composition for all samples grouped by the corresponding developmental stages. **d**. The assortativity analysis indicates an increased level of spatial clustering among similar cell types during heart development.

In order to quantitatively compare the change of spatial-temporal organization of cell type composition during embryonic heart development, we first examined the overall abundance of different cell types (**Fig. 2c**). We found that the abundance of ventricular cardiomyocytes increases dramatically during development (from an average of 25% in week 4.5-5 to 53% in week 9) (**Fig. 2c**). Notably, the abundance of atrial cardiomyocytes does not show this trend, which probably reflects the atrium compartments expand less dramatically compared to the ventricle compartments. Next, we compared the spatial organization patterns across developmental stages. Normal heart function relies on the coordinated activity of billions of cardiac cells; therefore we were interested to test whether spatially neighboring cells tend to belong to the same cell type. This is quantified by using a metric called the assortativity coefficient^21^, which is commonly used in social network analysis to characterize the tendency of friendship formed by similar individuals. In the current context, we considered the spatial network connecting neighboring cells. We further modified the definition of assortativity coefficient in order to account for the cellular heterogeneity within each spot location (see Methods for details). We found that the assortativity coefficient increased from 0.23 at week 4.5-5 to 0.50 at week 9 (Fig. 3d), suggesting the spatial organization becomes increasingly spatially coherent during heart development.

In conclusion, spatialDWLS is an accurate and computationally efficient method for estimating the spatial distribution of cell types from spatial transcriptomic data. Thus it provides a valuable enabling toolkit for investigating cell-cell interactions from various spatial transcriptomic technology platforms that do not have single-cell resolution. The spatialDWLS method can be easily accessed in Giotto ^17^, which is a user-friendly software package containing a large number of computational tools for spatial transcriptomic data analysis and visualization.

## Methods

### Cell type selection of spatial expression data by enrichment analysis

We use an enrichment based weighted least squares approach for deconvolution of spatial expression datasets. First, enrichment analysis using Parametric Analysis of Gene Set Enrichment (PAGE) method^22^ is applied on spatial expression dataset as previously reported^17^. The marker genes can be identified via differential expression gene analysis of Giotto based on the single cell RNA-seq data provided by users. Alternatively, users can also provide marker gene expression for each cell type for deconvolution. The number of cell-type specific marker genes is denoted by *m*. For each gene, we calculate the fold change as the ratio between its expression value at each spot and the mean expression of all spots. The mean and standard deviation of the fold change values are defined as μ and δ, respectively. In addition, we calculate the mean fold change of the *m* marker genes, which is defined as *S_m_*. The enrichment score (ES) is defined as follows:

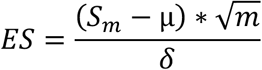

Then, we binarize the enrichment matrix with the cutoff value of ES = 2 to select cell types that are likely to be present at each point.

### Estimating cell type composition by using a weighted least squares approach

In previous work, we developed dampened weighted least squares (DWLS) ^9^ for deconvolution of single-cell RNAseq data. This method is extended here to deconvolve spatial transcriptomic data using the signature gene identification step described above. In short, DWLS uses a weighted least squares approach to infer cell-type composition, where the weight is selected to minimize the overall relative error rate. In addition, a damping constant *d* is used to enhance numerical stability, whose value is determined by using a cross-validation procedure. Here, we use the same sets of weights and damping constant across spots within same clusters to reduce technical variation. Finally, since the number of cells present at each spot is generally small, we perform another round deconvolution by remove those cell types that are predicted to present at a low frequency by imposing an additional thresholding (min frequency = 0.02 by default).

### Coarse-grained spatial transcriptomic data for model performance evaluation

The somatosensory cortex seqFISH+ data were abstained from https://github.com/CaiGroup/seqFISH-PLUS. To simulate spot-like data, we defined the square with 500 pixels time 500 pixels (~51.5μm) as one spot-like region. Then, average expression level was calculated for each spot-like region. Due to the small sample size, we only considered the 6 major clusters: excitatory neurons (eNeuron), inhibitory neurons (iNeuron), astrocytes, oligodendrocytes (Olig), microglia cells, and endothelial-mural cells (endo_mural).

### Benchmark comparison among different methods

Coarse-grained seqFISH+ dataset was used for benchmarking the accuracy of different deconvolution methods, including spatialDWLS, MuSiC, SPOTlight, stereoscope, and RCTD. Cell-type annotations for the original, single-cell resolution data were used as the ground-truth. All five methods use the same single cell RNA-seq dataset as a reference in deconvolution.

For spatialDWLS, we clustered the spot-like regions by using Leiden clustering as implemented in Giotto (Version 1.0.3) by using the following commands *createNearestNetwork(dimensions_to_use = 1:10, k = 4*) and *doLeidenCluster(resolution = 0.4, n_iterations = 1000)*

Then, marker genes of major clusters were identified by using the *findMarkers_one_vs_all* function with parameter setting: *method = ‘gini’, expression_values = ‘normalized’*. Top 100 ranked genes for each cell type were selected as marker genes. Average marker gene expression was calculated based on the cell type annotation of scRNA-seq. Then, deconvolution was applied by using the *runSpatialDWLS* function.

MuSiC ^8^ (version 0.1.1) was used for deconvolution by using whole single cell RNA-seq matrix. *ExpressionSet* classes were generated for both single cell RNA-seq (*SC.eset*) and spatial expression datasets (*ST.eset*). Then, cell type proportion was estimated by using *music_prop(bulk.eset = ST.eset, sc.eset = SC.eset)*

Then, to perform deconvolution by using SPOTlight ^15^ (version 0.1.0), signature genes were identified based on the major cell type annotation by using *Seurat::FindAllMarkers(logfc.threshold = 1, min.pct = 0.9)*.

Deconvolution was performed by using

*spotlight_deconvolution(se_sc = SC, counts_spatial = ST, cluster_markers = cluster_markers_all, clust_vr = “label”)*.

Next, we used stereoscope ^16^ (version 0.2.0) for the deconvolution of simulated dataset. Deconvolution was performed with parameter:

*stereoscope run -scc SC.tsv -scl cell_labels.tsv -stc ST.tsv -sce 5000.*

Finally, we used RCTD ^14^ (version 1.1.0) to evaluate the cell type composition for simulated seqFISH+ dataset. Signature genes were identified by using “*dgeToSeurat*”, then *“create.RCTD”* and “run.RCTD” were used to decompose the cell type composition. Finally, cell type percentage for each spot was calculated using the *“sweep”* function.

The computational efficiency of different methods was benchmarked by using the Visium brain dataset. All analyses were done on the same computer, which had Intel Xeon CPU E5-2650 2.00GHz and 380Gb memory. Of note, the Visium data cannot be used to evaluate accuracy because the ground-truth is not known.

### Root Mean Square Error (RMSE) calculation

Based on the cell type annotation of seqFISH+ dataset, we calculated the true cell type percentage for simulated spatial expression datasets. For a specific cell type, we divided spot-like regions into two groups based on the presence or absence of this cell type. RMSEs were calculated separately for these two groups.

### Analysis of a spatial transcriptomic dataset from the mouse brain

The Visium dataset was obtained from the 10X Genomics website (https://support.10xgenomics.com/spatial-gene-expression/datasets/1.1.0/V1_Adult_Mouse_Brain), which corresponds to a coronal section of the mouse brain. Then, Giotto was used for data analysis as (http://www.spatialgiotto.com/giotto.visium.brain.html). Only spots within tissue were kept for further analysis. Then, we filtered out low quality spots and genes by using filterGiotto with parameter:

*expression_threshold = 1, gene_det_in_min_cells = 50, min_det_genes_per_cell = 1000.*

After normalization and highly variable gene calculation, we performed neighborhood analysis with parameter: *createNearestNetwork (dimensions_to_use = 1:10, k = 15)* and clustered spots with parameter: *doLeidenCluster(resolution = 0.4, n_iterations = 1000)*. Finally, we used marker genes and scRNA-seq reported in Zeisel et al^18^ to deconvolute the Visium dataset.

### Analysis of a spatial transcriptomic dataset from developing mouse heart

The heart spatial transcriptomics datasets were obtained from ^20^. Then, we filtered out low quality spots and genes by using filterGiotto with parameter: *expression_threshold = 1, gene_det_in_min_cells = 10, min_det_genes_per_cell = 200*.

After normalization and highly variable gene calculation, we performed neighborhood analysis with parameter:

*createNearestNetwork(dimensions_to_use = 1:10, k = 10)*

and clustered spots with parameter: *doLeidenCluster(resolution = 0.4, n_iterations = 1000)*

In addition, we use the scRNA-seq data from the same website with spatial transcriptomics datasets. Based on the clusters reported, we re-analysed signature genes by using Giotto with parameter: *findMarkers_one_vs_all(method = ‘scran’)*

The average expression of marker genes was used for the deconvolution of heart ST datasets.

### Assortativity analysis

To evaluate the degree of spatial coherence, we extended the assortativity analysis^21^, a method commonly used in the network analysis to evaluate the tendency of similar networks nodes are connected to each other. Here, we generated a spatial network by connecting spots that are immediately next to each other. The associativity coefficient represents the normalized deviation of edges connecting the same cell type than expected by chance. More precisely, it is defined by the following formula:

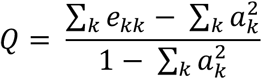

where

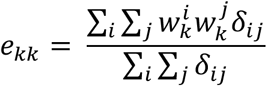

and

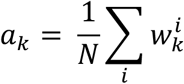

In the above, 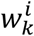 represents the fraction of cell-type *k* at the *i*-th spot, *N* represents the total number of spots, and *δ_ij_* is the Kronecker-delta defined as

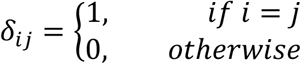

If the values of 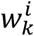 are binary, then the above definition reduces to the original formula in ^21^.

## Data and code availability

All codes, data, and analysis results in this paper are publicly available at GitHub: https://github.com/rdong08/spatialDWLS_dataset. Furthermore, the spatialDWLS method is implemented as the *runDWLSDeconv* function in Giotto (https://github.com/RubD/Giotto) and detailed tutorial and vignette are available at Giotto website (http://www.spatialgiotto.com). A vignette showing the use of spatialDWLS in Giotto can be found at Github (https://github.com/rdong08/spatialDWLS_dataset/blob/main/codes/Heart_ST_tutorial_spatialDWLS.Rmd).

## Acknowledgements

We gratefully acknowledge the helpful discussion with Dr. Ruben Dries. This research was supported by NIH grants UH3HL145609 and R01AG066028 to G.-C.Y.

## Supplementary Figure Legends

**Figure S1.**
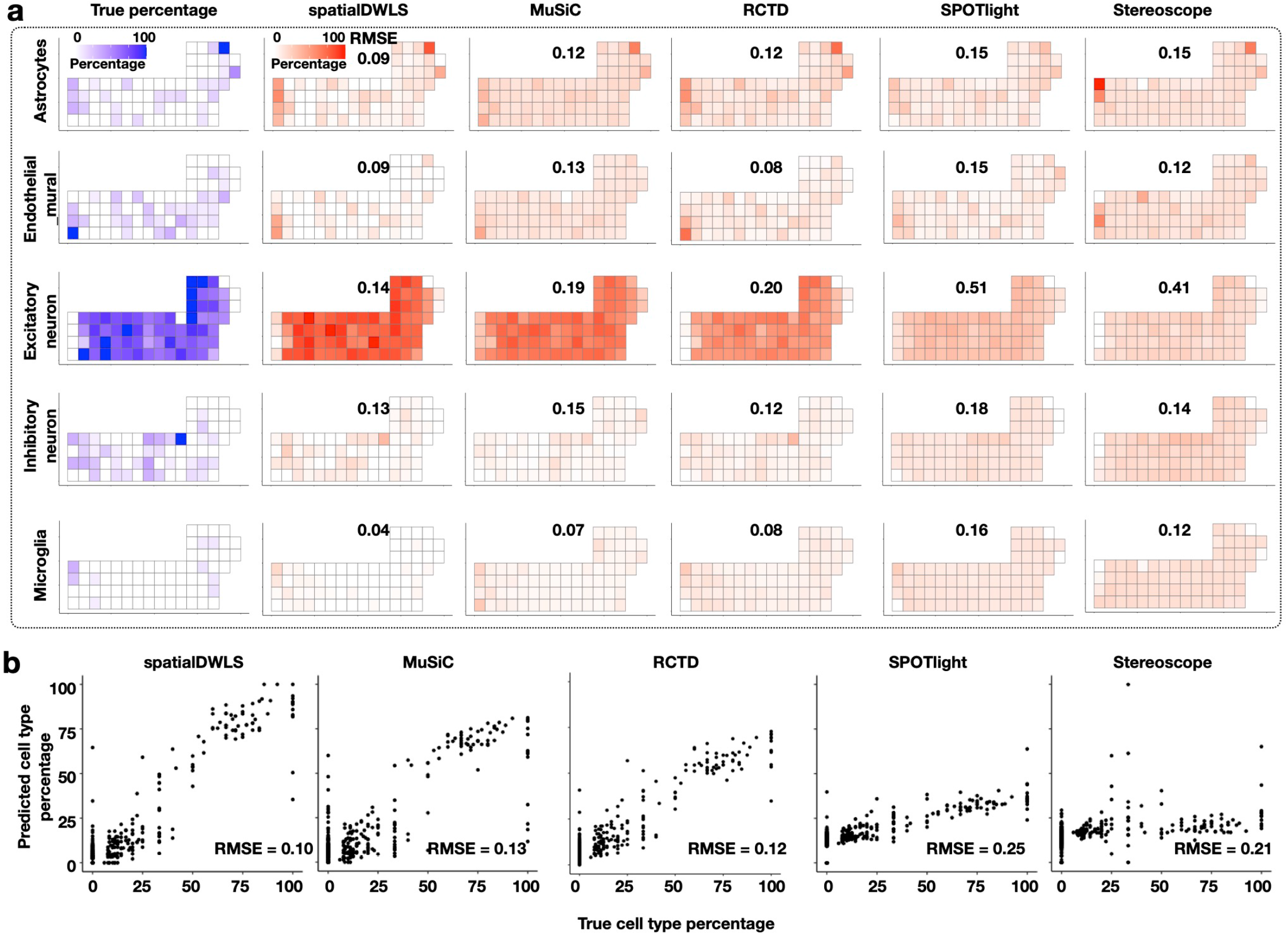
Extended comparison of the accuracy of different deconvolution methods (see Figure 1b for description). **a.** Heatmaps indicate the spatial distribution of different cell types. The true frequency (blue) are compared with estimates from different deconvolution methods (red). The RMSE corresponding to each cell type is indicated. **b.** Scatter plot indicate the relationship between true (y-axis) and inferred (x-axis) cell type frequencies. Results from all the cell types are overlaid. The overall level of RMSE is indicated for each method.

**Figure S2.**
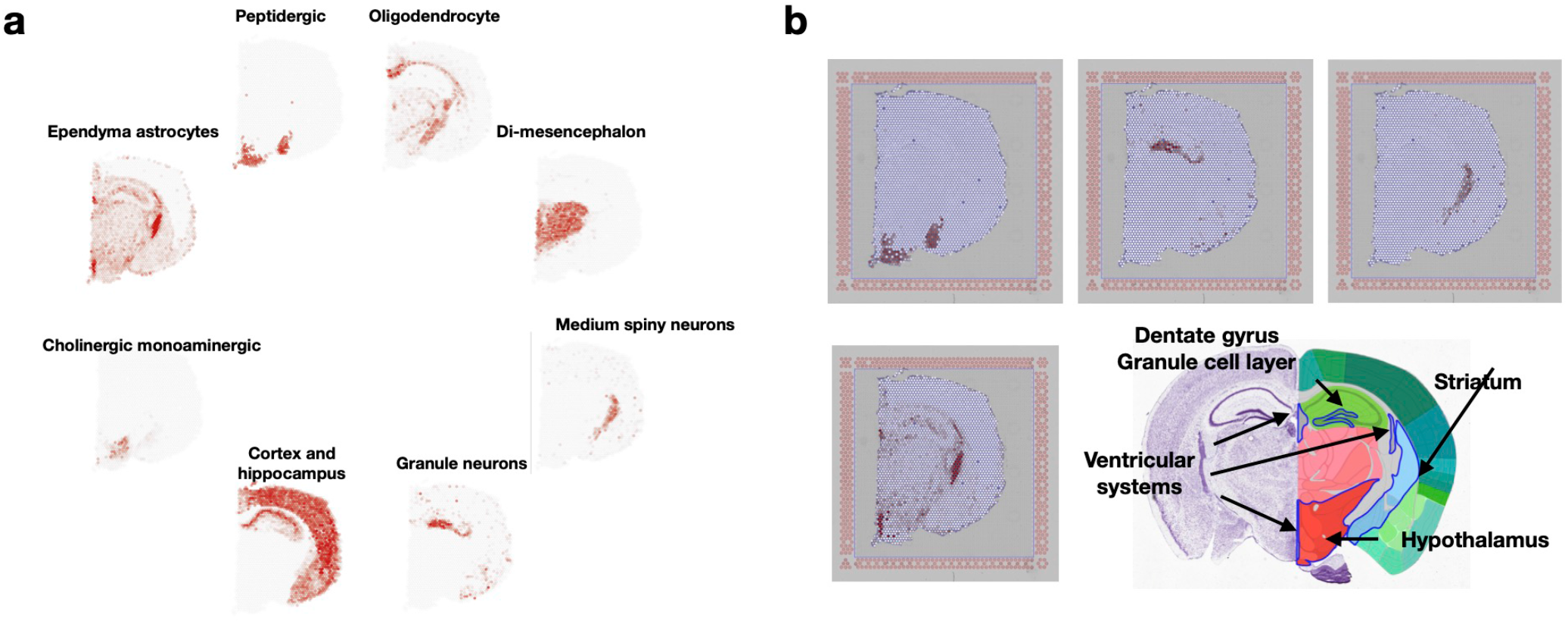
Deconvolution analysis of the mouse brain Visium dataset. **a.** The spatialDWLS estimated spatial distribution of different cell types (indicated as red dots). **b.** The anatomic structure of the brain, obtained from the Allen Brain Atlas, is shown here for comparison with the deconvolution analysis results.

**Figure S3.**
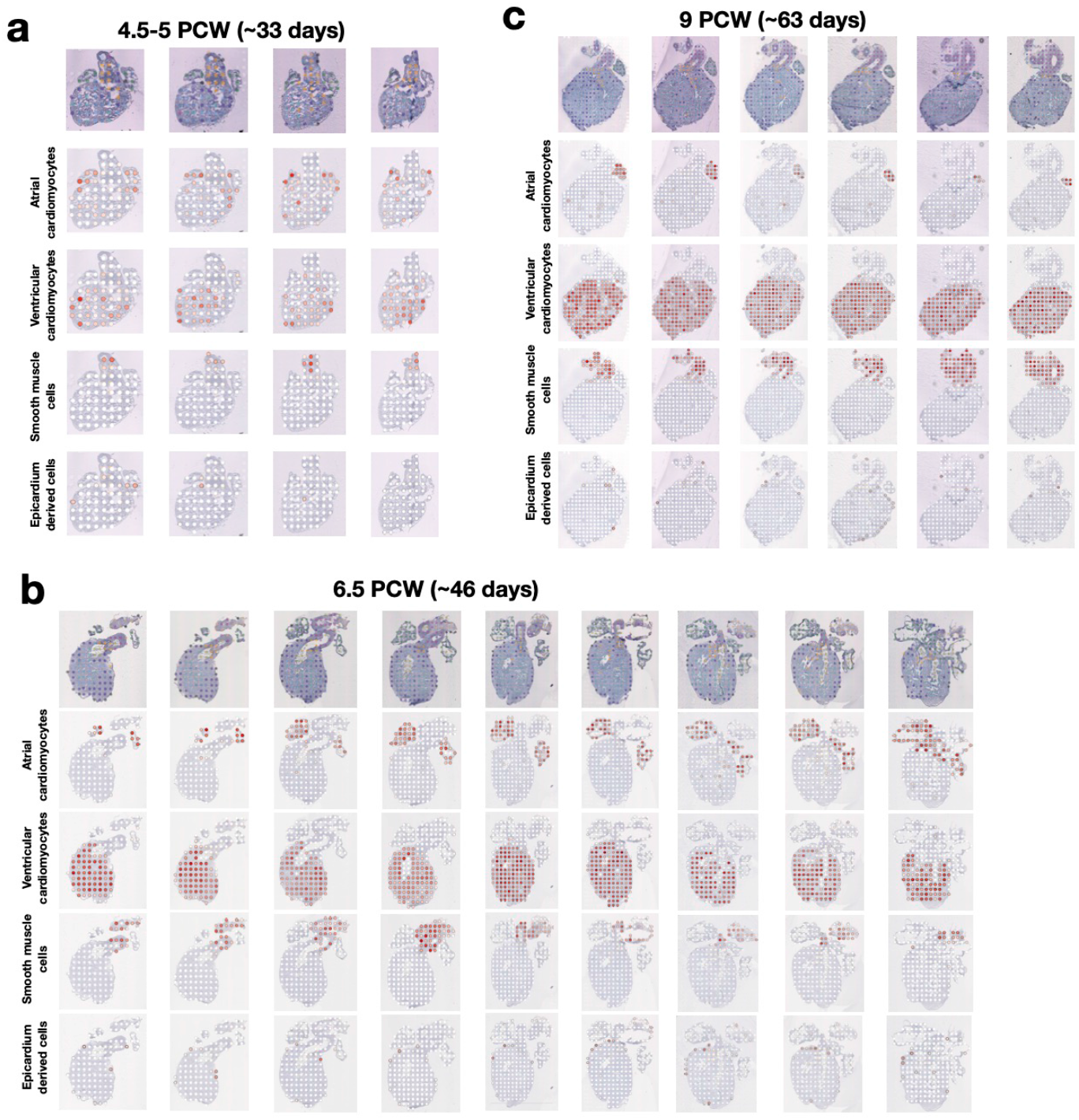
Deconvolution analysis of the human developing heart Spatial Transcriptomic dataset. SpatialDWLS estimated spatial distribution of atrial cardiomyocytes, ventricular cardiomyocytes, smooth muscle cells and epicardium derived cells in week 4.5-5 (**a**), week 6.5 (**b**) and week 9 (**c**). The magnitude of cell-type frequency at each spot is indicated by red circles.

